# Transcriptomic and network analyses of an alcohol-induced peripheral neuropathy model identify putative role for histone demethylase *Jmjd1c*

**DOI:** 10.1101/2025.10.06.680792

**Authors:** Walker D. Rogers, Lauren Moncayo, Zain Akbar, Madison Cruz, Abdel-Rahman Dahman, Ammar Mohiuddin, M Imad Damaj, Michael F. Miles

**Author notes:** **Corresponding authors:** Michael Miles, MD, PhD, and M. Imad Damaj, PhD **E-mail:**. Contributed equally to the manuscript (Walker D. Rogers, Lauren Moncayo).

## Abstract

**Background:** Alcohol-induced peripheral neuropathy (AIPN) is a painful and prevalent condition associated with chronic alcohol use, yet its molecular underpinnings remain poorly understood. Because the analgesic effects of ethanol may reinforce alcohol consumption, elucidating the mechanisms driving AIPN is essential. This study aimed to identify ethanol-regulated gene expression patterns in the nervous system of a mouse model of AIPN.

**Methods:** Male (n = 10) and female (n = 12) C57BL/6J mice were administered either an ethanol-containing Lieber-DeCarli liquid diet at 5% or an isocaloric control diet for four weeks. Ethanol consumption was recorded daily for the experimental group. After the drinking protocol, spinal cord and dorsal root ganglia tissues were collected for RNA sequencing.

**Results:** Ethanol-regulated genes were identified for each sex-tissue group using DESeq2, and results were compared to known rodent neuropathic pain gene signatures. Weighted gene co-expression network analysis (WGCNA) identified modules of co-expressed genes associated with ethanol administration. Hub genes with high intramodular connectivity were identified for ethanol-correlated modules. Of the 14 identified hub genes, 10 have been previously implicated in pain or neuropathy, including *Jmjd1c*, *Phf8*, and *Gas6*, which emerged as particularly strong candidates for involvement in AIPN pathophysiology.

**Conclusions:** These findings provide novel insights into the gene networks underlying AIPN and nominate specific genes for future functional studies.

## Introduction

Peripheral neuropathy (PN) is a condition in which neurons outside the central nervous system are damaged. Depending on whether sensory, motor, or autonomic nerves are affected, symptoms can include muscle weakness, painful irritation, numbness, or dysregulated thermal regulation (England & Asbury, 2004). PN has a multifactorial etiology, with diabetes, autoimmune disorders, exogenous toxin exposure, and other factors all potentially contributing to its development. Due to the multiple pathways noted here, PN is extremely widespread. Nearly 8% of people 18 and older estimated to have some form of PN, and prevalence increases dramatically with age (Visser et al., 2015). People with painful peripheral neuropathy report a decreased quality of life, with severity of pain significantly negatively correlated with overall quality of life (Girach et al., 2019). Beyond the direct effect of pain, PN has been associated with anxiety, depression, and sleep dysregulation (Naranjo et al., 2019). Because of the high prevalence in the population and the reduction in quality of life associated with PN, it is critical to understand the causal mechanisms underlying the development and persistence of peripheral nerve damage.

One common, though understudied, form of PN is alcohol-induced peripheral neuropathy (AIPN), which occurs in a substantial proportion of individuals with alcohol use disorder (AUD). Epidemiological studies estimate that over 46% of people with AUD also meet criteria for AIPN (Julian et al., 2019). The exact pathogenesis of AIPN is not fully understood but is thought to involve the direct neurotoxic effects of ethanol on peripheral neurons, mitochondrial dysfunction, and possibly nutritional deficiencies that are common in people with AUD (Julian et al., 2019). Histopathological studies have described length-dependent axonal degeneration (Wöhrle et al., 1998), demyelination (Tessitore et al., 2022), and sensory fiber loss (Kokotis et al., 2023), but the molecular mechanisms driving these changes have not been fully characterized.

The relationship between pain conditions and alcohol use is bidirectional and complex. Ethanol dose-dependently produces analgesic effects that can acutely reduce pain, potentially reinforcing continued alcohol consumption (Thompson et al., 2017). Indeed, the practice of self-medicating pain symptoms with alcohol use has previously been described. Around 25% of individuals with orofacial pain or arthritis reported using alcohol to manage their pain symptoms in a community-based survey (Riley & King, 2009), and 38% of primary care patients with a history of illicit drug use and recent heavy alcohol use reported drinking to cope with pain (Alford et al., 2016). However, in the case of alcohol-induced peripheral neuropathy (AIPN), alcohol use itself serves as the initiating factor for the painful condition. This creates a maladaptive feedback loop in which alcohol-induced nerve damage promotes pain, which may in turn encourage further drinking. Intervening in this cycle requires a better understanding of how chronic ethanol exposure alters the molecular landscape of the peripheral and central nervous systems.

Transcriptomic profiling is one method for uncovering the molecular pathways associated with disease phenotypes. RNA sequencing enables the detection of differentially expressed genes, pathway perturbations, and regulatory elements relevant to neuropathic pain. However, existing peripheral neuropathy RNA-seq studies largely focus on surgical and metabolic models of PN (Bolívar et al., 2024; Hinder et al., 2018; Welleford et al., 2020) or neuropathy induced by other toxins (Li et al., 2021; Naratadam et al., 2024). Furthermore, studies often focus on a single sex or tissue type, potentially overlooking important biological variation.

This study aims to address these gaps by performing a transcriptomic analysis of dorsal root ganglia (DRG) and spinal cord (SC) tissues in both male and female mice exposed to chronic high-dose ethanol via the Lieber-DeCarli liquid diet. These tissues were chosen for their relevance to nociception: the DRG houses primary sensory neurons, while the spinal cord serves as a critical relay and integration center for pain signaling. Using differential gene expression analysis and weighted gene co-expression network analysis (WGCNA), we identify gene expression changes and co-regulated gene networks associated with ethanol exposure. This approach enables the identification of candidate genes and pathways involved in the development and maintenance of AIPN, advancing our understanding of its molecular basis and pointing to potential therapeutic targets.

## Materials and Methods

### Animals

Seven-week old adult male (n = 14) and female (n = 12) C57BL/6J mice were purchased from Jackson Laboratory (Bar Harbor, ME). Mice were housed in pairs in an AAALAC-accredited animal care facility on a regular 12-hour light/dark cycle with food and water provided *ad libitum* (7012 Teklad LM-485 Mouse/Rat Sterilizable Diet, Harlan Laboratories Inc., Indianapolis, IN). Animals were given a one-week acclimation period before initiating the alcohol liquid diet drinking study for four weeks. After the experimental protocol, mice were euthanized by decapitation following protocols approved by the Virginia Commonwealth University Institutional Animal Care and Use Committee. Animal welfare was monitored daily during the study.

### Alcohol

A 5% (v/v) ethanol solution was prepared from 100% Biology Grade Ethanol (Sigma-Aldrich, St. Louis, MO) diluted in sterile deionized water.

### Lieber-DeCarli Drinking Protocol

The Lieber-DeCarli control (F1259) and EtOH (F1258) liquid diet was purchased from Bio-Serv (Flemington, NJ) and prepared similarly to previously published methods (De Logu et al., 2019). The control liquid diet was calorically matched to ethanol liquid diet with maltose-dextrin to isolate the physiological effects of the ethanol administration.

Each pair-housed cage of mice was transitioned from regular chow and water to 50 mL of control liquid diet for one week of habituation. After one week habituating to the control liquid diet, mice were single housed and given 30 mL of control or 5% (v/v) EtOH liquid diet. Following this habituation, mice were single housed and given 30 mL of either control or 5% (v/v) ethanol-containing liquid diet. Every 24 hours, the volume of liquid diet consumed was recorded, mouse body mass was measured, and fresh diet was provided. Mice were maintained on their assigned diets for four weeks prior to tissue collection. Recently, our group has shown that these ethanol concentration and duration parameters have been shown to decrease caudal nerve conduction amplitude and intraepidermal nerve fiber density in a mouse model of AIPN, and thus we used the same experimental design for the present study (Moncayo et al., 2025).

### Tissue Collection and RNA Isolation

For tissue-specific transcriptomic analyses, spinal cord (overlaying the T13-L1 vertebrae) and dorsal root ganglia (L4-L6 lumbar DRG) were collected and flash frozen in liquid nitrogen within five minutes of euthanasia. Total RNA was extracted from tissue samples using the Qiagen miRNeasy Mini Kit (Cat. No. 217004) following the manufacturer’s protocol. Tissue was homogenized in STAT60 (Amsbio, UK). RNA quality control was performed using a NanoDrop spectrophotometer and Agilent Bioanalyzer, and all samples yielded a 260/280 ratio ∼ 2.0 and an RNA Integrity Number ≥ 8.0.

### RNA-seq

Sequencing was conducted at the VCU Genomics Core on an Illumina NextSeq 2000 with P3 flow cell (Illumina, Inc. San Diego, California). Read quality control, adapter trimming, duplicate identification, and GC content analysis were conducted using fastp (v0.23.2) with default settings (Chen et al., 2018). Read strandedness was verified with default parameters in the how_are_we_stranded_here Python library (v1.0.1) (Signal & Kahlke, 2022). Reads that passed quality control were aligned to the GRCm39 genome by STAR (v2.7.10b) (Dobin et al., 2013). BAM files were sorted and indexed by Samtools (v1.5.1) (Danecek et al., 2021). Read counting was performed using the featureCounts function from the Subread package (v2.0.1) (Liao et al., 2013). Principal component biplots of variance-stabilized transformed reads were generated to detect outliers. A total of 35 RNA-seq samples in the following experimental groups passed quality control and were further analyzed: 8 male DRG samples (n = 4 ethanol, n = 4 control), 10 male SC samples (n = 5 ethanol, n = 5 control), 8 female DRG samples (n = 4 ethanol, n = 4 control), and 9 female SC samples (n = 5 ethanol, n = 4 control).

### Statistical Analysis

Differential gene expression analysis was performed using DESeq2 (v1.40.2) (Love et al., 2014). Differentially expressed genes between ethanol treatment groups were defined as having an adjusted p-value < 0.1. Data visualization was conducted using ggplot2 (v3.5.1) (Wickham, 2016). Gene set overlap was assessed via Fisher’s exact test using the GeneOverlap package (v1.44.0) (Shen, 2025). Genes with fewer than 10 raw counts in at least four (the smallest experimental group size) samples were filtered out prior to DEG analysis and WGCNA. Background sets for overlap comparisons were based on the number of post-filter transcripts during RNA-seq, and the sets were tissue- and sex-specific: 16,795 male DRG genes, 17,123 male spinal cord genes, 16,048 female DRG genes, and 16,564 female spinal cord genes.

Gene ontology and KEGG pathway enrichment analysis were conducted with the enrichGO and enrichKEGG functions, respectively, from the clusterProfiler R package (Yu et al., 2012). Ontology categories and pathways were considered enriched if they yielded an adjusted p-value < 0.05.

A comparison with an existing neuropathic pain transcriptional signature was conducted using the overlap between this study’s sex- and tissue-specific ethanol DEGs and the list of common neuropathic pain genes from DRG found in Table S2 in Pokhilko et al. (Pokhilko et al., 2020). The GeneOverlap package mentioned previously was used to perform a Fisher’s exact test for each overlap comparison.

Weighted gene co-expression network analysis (WGCNA) was conducted for each sex-tissue group using the WGCNA package (v1.73) (Langfelder & Horvath, 2008). Analysis was performed on all filtered transcripts as referenced above. Parameters included: unsigned network type, minimum module size of 100, deep split = 1, and dendrogram cut height = 0.25. Soft-thresholding powers were: 10 for male DRG, 6 for male spinal cord, 9 for female DRG, and 8 for female spinal cord. WGCNA modules were tested for enrichment of ethanol DEGs from corresponding sex-tissue groups using the Fisher’s exact test implemented in the GeneOverlap package. Hub genes were identified using the chooseTopHubInEachModule function from WGCNA, which uses intramodular connectivity (K_IM_) as the hub gene selection criteria. Ethanol-associated gene modules were identified by calculating Pearson’s correlations between module eigengenes and ethanol administration variables. Modules with a raw p-value < 0.05 for the eigengene-ethanol correlation were considered significantly correlated.

## Results

### Ethanol Intake in Male and Female Mice on Lieber-DeCarli Diet

Daily ethanol intake (g/kg) was measured over 28 days in male and female mice administered the Lieber-DeCarli liquid diet with 5% (v/v) ethanol (**Figure 1A**). Female mice consumed an average of 18.2g/kg EtOH daily, and male mice consumed 17.6g/kg EtOH daily. Female mice consistently consumed ethanol at relatively stable levels, with average daily intakes ranging from 14.3-23.9 g/kg across the study. Male mice exhibited more variable ethanol intake over the 28 days, with average daily consumption ranging from 9.5-27.5 g/kg, though average intakes largely overlapped between sexes.

**Figure 1.**
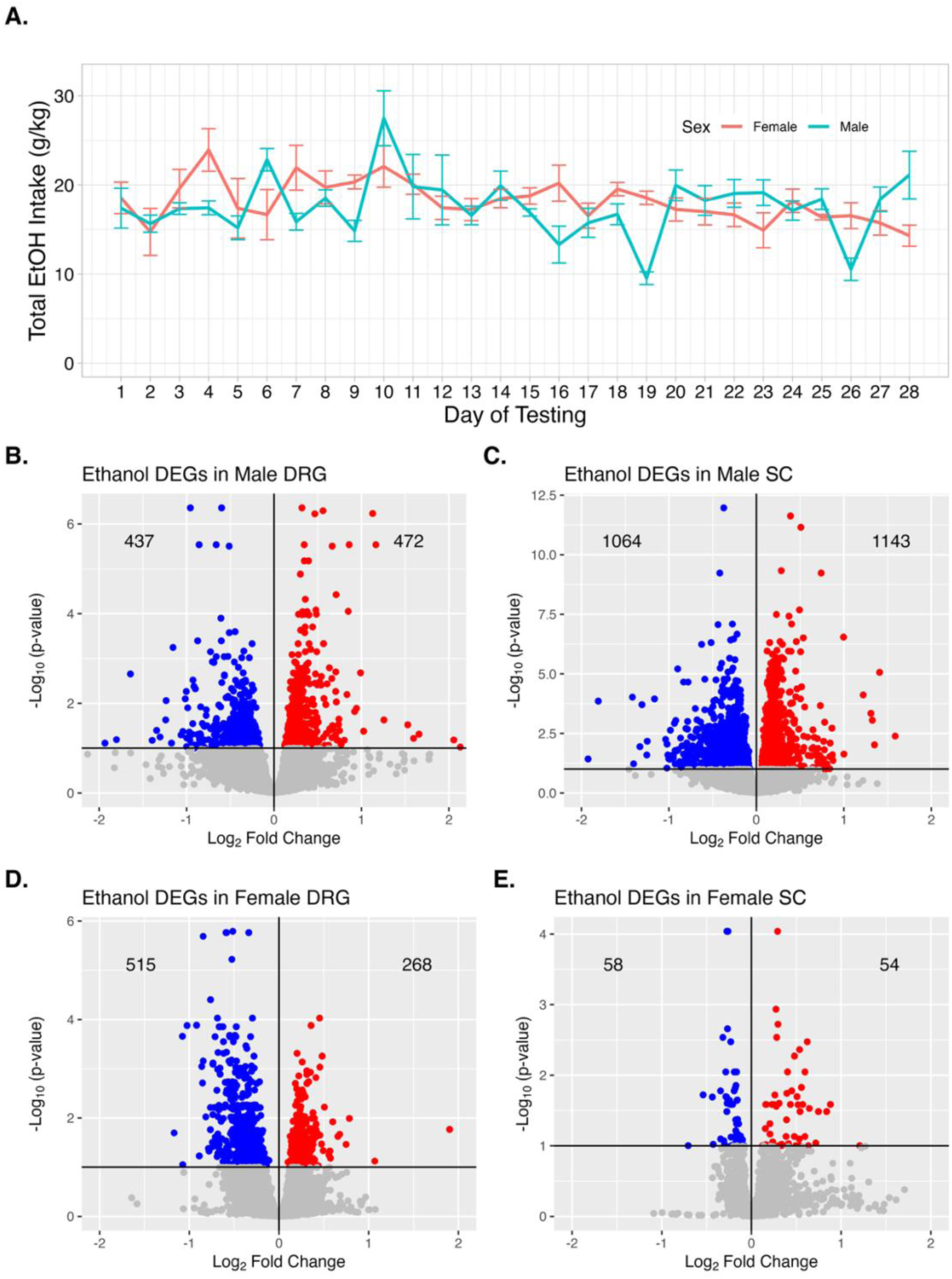
A: Average daily ethanol intake for male and female mice on 5% ethanol Lieber-DeCarli liquid diet. Error bars represent +/-1 standard error. B-E: Volcano plots of genes regulated by chronic ethanol exposure. Horizontal line indicates significance cutoff of p_adjust_ < 0.10. The number of upregulated and downregulated genes is included in each panel.

### Identification of Differentially Expressed Genes

Differential expression analysis using DESeq2 revealed substantial transcriptional changes in response to chronic ethanol consumption. In male mice, 909 ethanol-regulated genes were identified in dorsal root ganglia (DRG) and 2,207 in spinal cord tissue (**Figure 1B-C; Table S1A-B**). Of the DRG DEGs, 437 were downregulated and 472 were upregulated. In the spinal cord, 1,064 genes were downregulated and 1,143 were upregulated. There was significant overlap between the male DRG and spinal cord DEG sets, with 348 shared genes (Fisher’s exact test: OR = 5.07, *p* < 0.0001; **Table S1E**). In female mice, 783 DEGs were identified in DRG (515 downregulated, 268 upregulated), and 112 in spinal cord (58 downregulated, 54 upregulated) (**Figure 1D-E**; **Table S1C-D**). Although the overall number of DEGs was lower in the spinal cord, 15 genes were shared between the two tissues (OR = 3.18, *p* < 0.001; **Table S1F**). Gene set overlaps are visualized in **Figure S1**.

### Overrepresentation Analysis

Gene ontology (GO) and KEGG pathway enrichment analysis was conducted to characterize the biological systems modulated by chronic high-dose ethanol and potentially underlying alcohol-induced peripheral neuropathy. In DRG from male mice, the 909 ethanol DEGs were enriched for 348 unique GO/KEGG terms (**Table S2A**) across the aspects of biological process, molecular function, cellular component, and functional pathway. Neuron- and muscle-associated terms were highly represented (**Figure 2A**), as were potassium, sodium, and calcium ion transport and neuron apoptotic processes. The set of 2207 spinal cord ethanol DEGs derived from male samples was enriched for 1470 GO/KEGG terms (**Table S2B**). As seen in DRG results, GO terms associated with monoatomic ion transport and neuron death were highly significant in the spinal cord gene set (**Figure 2B**). Additionally, genes for the development, myelination, and axon ensheathment of the peripheral nervous system were all significantly overrepresented in male spinal cord ethanol DEGs.

**Figure 2.**
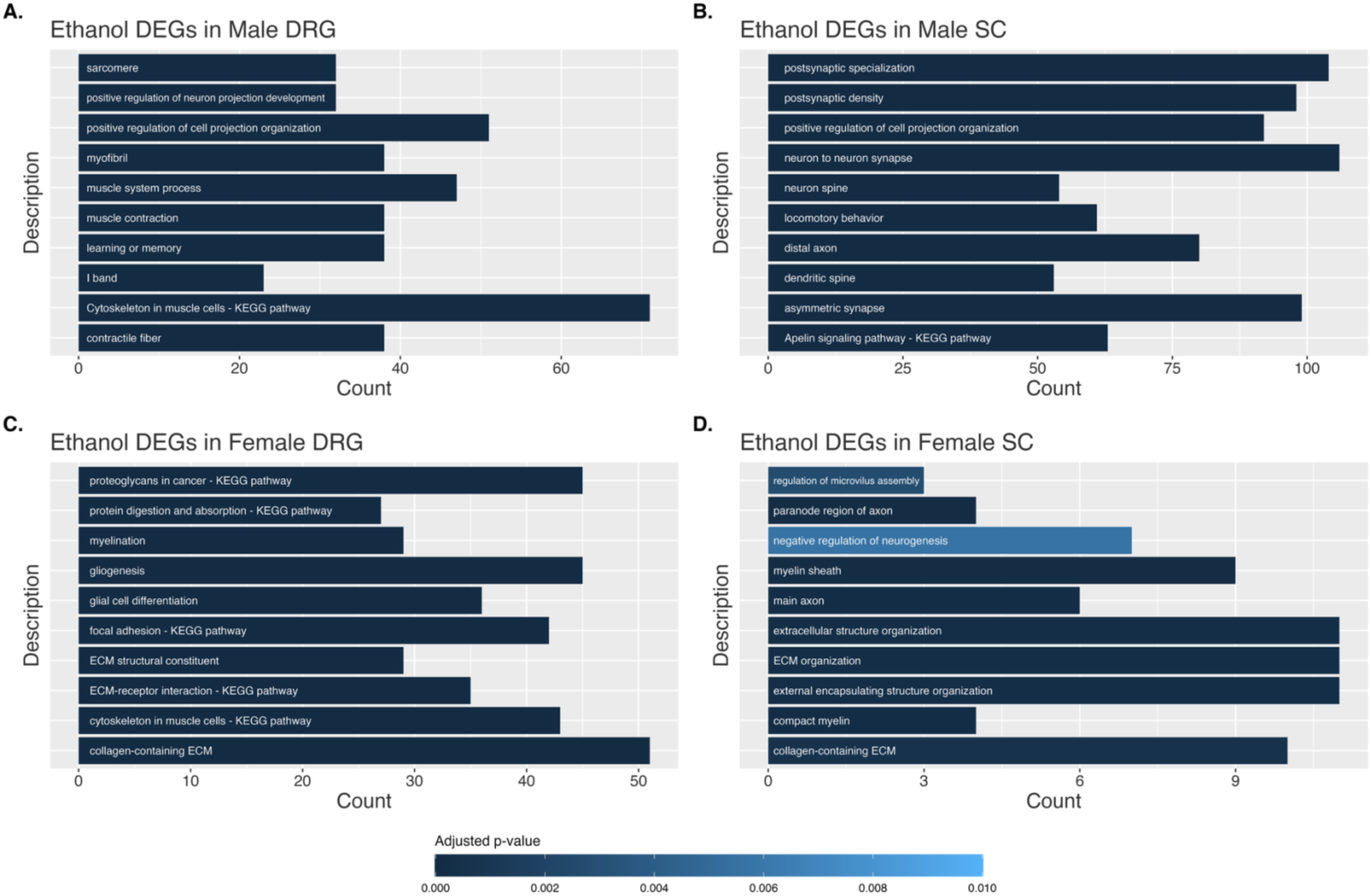
The 10 GO terms and KEGG pathways most significantly enriched in each set of ethanol differentially expressed genes for each sex-tissue group. KEGG = Kyoto Encyclopedia of Genes and Genomes.

In DRG DEGs from female mice, the most significantly enriched gene ontology and KEGG categories again included myelination and peripheral nervous system development, as well as terms related to the structure and function of the extracellular matrix (**Figure 2C**; **Table S2C**). Female spinal cord-derived ethanol DEGs were enriched for genes regulating myelin (myelin sheath, compact myelin), the extracellular matrix (ECM organization, collagen-containing ECM), and neurons (negative regulation of neurogenesis, main axon, response to axon injury) (**Figure 2D**; **Table S2D**).

### Relationship with Neuropathic Pain Signatures

Gene sets derived from rodent nociception and neuropathy studies were queried for overlap with ethanol DEG sets from the current analysis to identify similar transcriptomic profiles across pain conditions. In male mice, DRG DEGs overlapped significantly with the rodent peripheral neuropathy signature from an analysis of multiple RNA-seq datasets from DRG by Pokhilko et al. (Pokhilko et al., 2020) (OR = 3.29, p < 0.0001) (**Table S3A-B**). Genes present in this overlap include *Oprd1*, *Cacna2d1*, and *Kcnk1*.

Ethanol DEGs in spinal cord from male mice did not significantly overlap the Pokhilko et al. gene set (OR = 1.26, p = 0.12) (**Table S3C**). DEG sets from female DRG and spinal cord samples similarly yielded non-significant overlaps (**Table S3D-E**).

### Weighted Gene Co-expression Network Analysis

Modules of highly co-expressed genes were identified separately for each sex and tissue type to elucidate potential druggable targets associated with chronic alcohol-induced transcriptomic changes (see **Table S6** for module membership). Pearson correlations between module eigengenes and a binary ethanol administration variable were calculated, and GO analysis was performed on modules with a significant uncorrected correlation p-value. Variable-eigengene correlations were also performed on the continuous ethanol drinking measurements of average daily consumption and total ethanol consumption, and these results showed a high degree of concordance with the binary ethanol administration variable (**Figure S2**). Significant module-ethanol associations were found in all sex-tissue analyses (**Figure 3**). Of the 14 significant modules, 12 yielded significant GO enrichment analysis results (**Table S4**). The most highly enriched GO terms for these sex- and tissue-specific results are summarized in **Table 1**, which highlights shared and distinct biological pathways implicated in chronic ethanol exposure. As a complementary analysis, all WGCNA modules were queried for enrichment of ethanol DEGs from their respective sex-tissue group. Ten WGCNA modules were enriched for ethanol DEGs according to an FDR-corrected p-value < 0.05 (**Table 2**, full overrepresentation results in **Table S5**).

**Figure 3.**
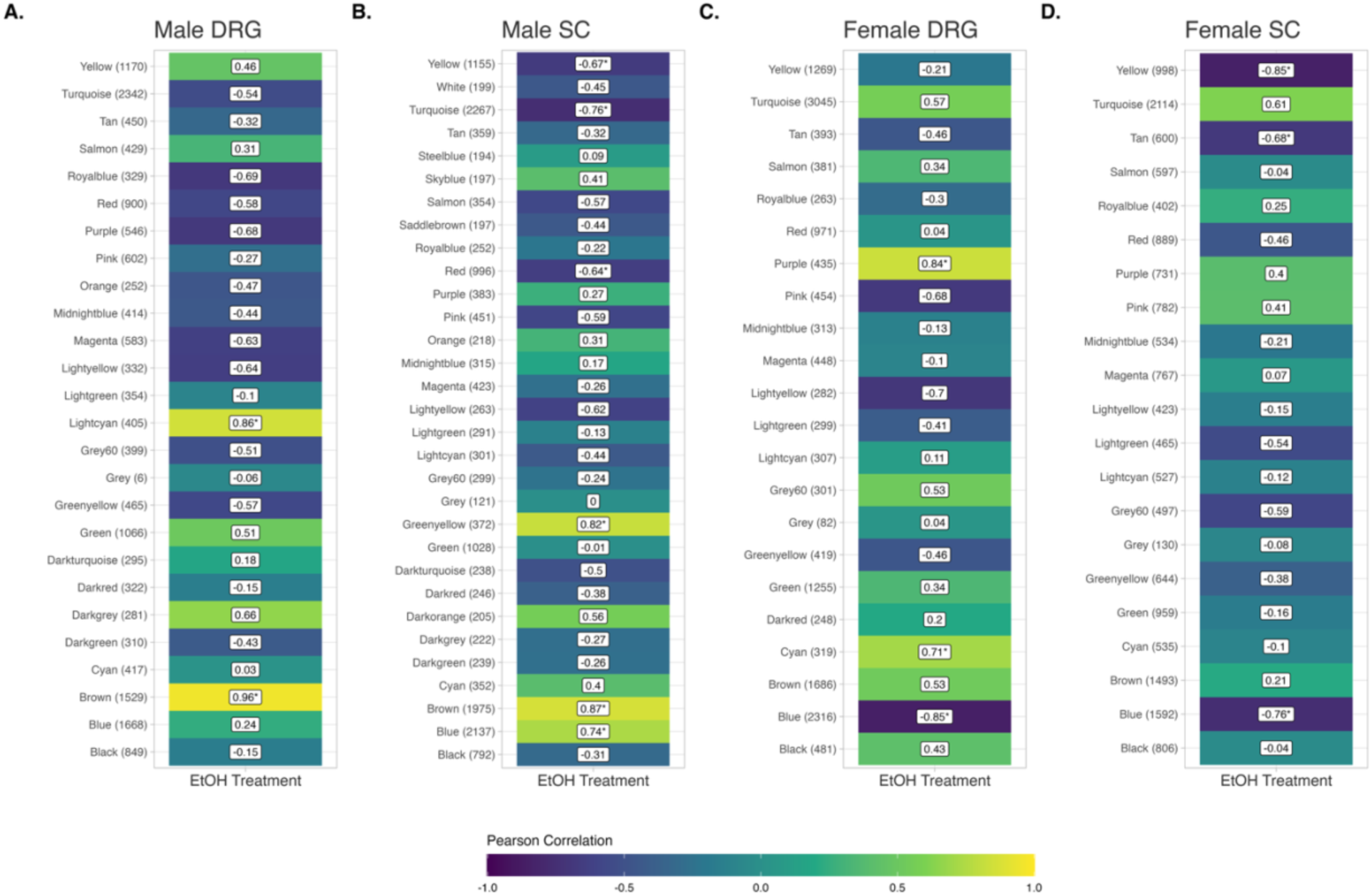
Heatmaps of Pearson correlation coefficients between WGCNA module eigengenes and binary ethanol administration variables. Correlations significant at an uncorrected p-value of 0.05 are denoted with an asterisk. The number of genes in each module is indicated in parentheses. DRG = dorsal root ganglia; SC = spinal cord.

**Table 1.** Ethanol-associated WGCNA modules with corresponding enriched GO terms for each sex-tissue group. The three most significantly enriched GO terms are represented, unless fewer than three ontologies are enriched (i.e., Female DRG Cyan module). An FDR-corrected p-value < 0.10 was used for the enrichment significance threshold. Full GO results for ethanol-regulated genes are reported in Table S2. DRG = dorsal root ganglia; ECM = extracellular matrix; NAD = nicotinamide adenine dinucleotide; PNS = peripheral nervous system.

| **Sex** | **Tissue** | **Module** | **Selected Enriched GO Terms** |
| --- | --- | --- | --- |
| Male | DRG | Brown | Inclusion body, myelin sheath, DNR replication preinitiation complex |
| Male | SC | Brown | Distal axon, postsynaptic density, asymmetric synapse |
|  |  | Turquoise | Cytosolic ribosome, ribosome, cytoplasmic translation |
|  |  | Blue | Asymmetric synapse, postsynaptic specialization, postsynaptic density |
|  |  | Yellow | phosphatidylinositol binding, phosphatidylinositol bisphosphate binding, acyltransferase activity |
|  |  | Red | collagen-containing ECM, ECM structural constituent, myelin sheath |
| Female | DRG | Blue | Schwann cell differentiation, PNS development, gliogenesis |
|  |  | Cyan | Microtubule |
|  |  | Purple | ATP-dependent protein folding chaperone, unfolded protein binding, protein folding chaperone |
| Female | SC | Blue | negative regulation of axonogenesis, negative regulation of neurogenesis, cognition |
|  |  | Yellow | Histone deacetylase complex, myelin sheath, NAD metabolic process |
|  |  | Tan | mitochondrial protein-containing complex, mitochondrial respirasome, respiratory chain complex |

**Table 2.**
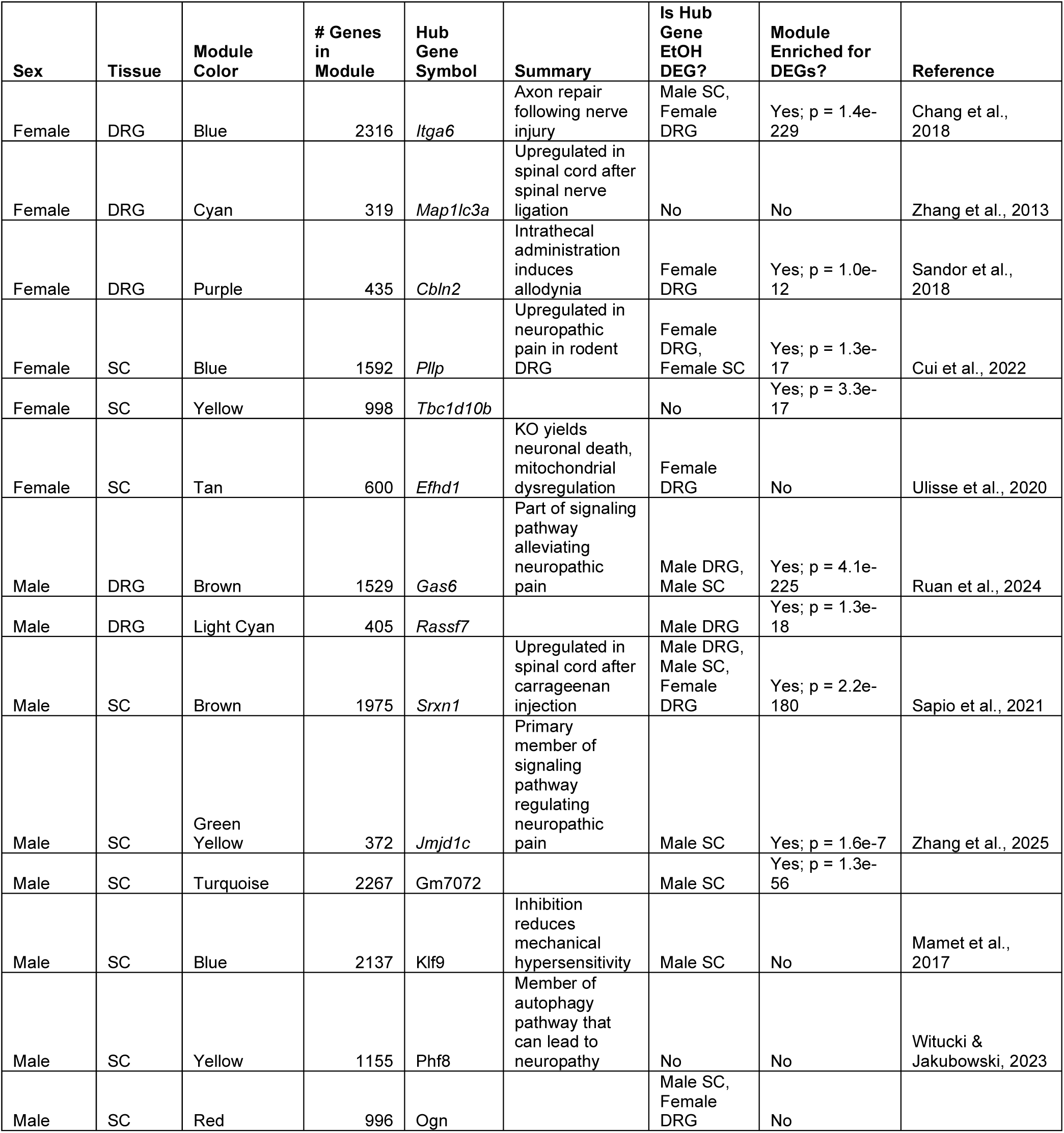
Hub genes with the highest intramodular connectivity for WGCNA modules significantly correlated with ethanol treatment. DRG = dorsal root ganglia; KO = knock-out.

Within each ethanol-associated module, the top hub gene, defined by highest intramodular connectivity, was identified as a potential regulator of module-wide expression patterns (**Table 2**). Ten of the fourteen hub genes (*Itga6*, *Map1lc3a*, *Cbln2*, *Pllp*, *Tbc1d10b*, *Efhd1*, *Gas6*, *Rassf7*, *Jmjd1c*, *Phf8*) were previously implicated in pain-or neuropathy-related studies, supporting their relevance to AIPN pathology.

## Discussion

Chronic alcohol use damages both small and large fibers of the peripheral nervous system, contributing to alcohol-induced peripheral neuropathy (AIPN), a common and debilitating complication of alcohol use disorder (AUD). Given that neuropathic pain may negatively reinforce alcohol consumption through ethanol’s analgesic effects, preventing AIPN could reduce the disease burden associated with AUD and improve long-term outcomes. Understanding the molecular genetic mechanisms driving AIPN is therefore crucial to therapeutic development. This study presents a transcriptomic analysis of a mouse model of high-dose ethanol exposure, reflecting heavy drinking in humans with AUD, that reproduces behavioral pathology seen with AIPN (Moncayo et al., 2025).

Previous studies have identified transcriptomic changes in rodent DRG following chemotherapeutic administration (Naratadam et al., 2024) and nerve injury (Barry et al., 2023), but to our knowledge, no studies have yet examined AIPN-specific transcriptional patterns. We identified ethanol-regulated genes and gene co-expression networks in the dorsal root ganglia (DRG) and spinal cord of male and female mice that have mechanistic implications on AIPN. Notably, multiple central hub genes within these networks are validated in prior studies on pain mechanisms and may represent viable targets for therapeutic intervention. Our results also expand on prior literature by providing a sex-stratified analysis of ethanol-induced gene expression changes relevant to neuropathic pain.

Previous studies have identified transcriptomic changes in rodent DRG following chemotherapeutic administration (Naratadam et al., 2024) and nerve injury (Barry et al., 2023), but to our knowledge, no studies have yet examined AIPN-specific transcriptional patterns. Here, we characterized gene expression changes in DRG and spinal cord tissues after chronic high-dose ethanol exposure via the Lieber-DeCarli liquid diet.

The liability to develop peripheral neuropathy is influenced by both genetic and environmental factors. While inherited neuropathies are typically driven by pathogenic variants in specific genes, some of these same genes may also be regulated by chronic ethanol exposure. For example, *Mpz* encodes myelin protein zero, a key structural component of the myelin sheath in peripheral nerves. Variants in *MPZ* are among the most common causes of the autosomal dominant neuropathy Charcot-Marie-Tooth disease Type 1 (Sanmaneechai et al., 2015), and *Mpz* was significantly downregulated in both male spinal cord and female DRG samples in our study. Ethanol-induced reductions in *Mpz* expression may therefore contribute to demyelination and axonal dysfunction in AIPN. Similarly, *Pmp22* (peripheral myelin protein 22), which maintains myelin integrity in Schwann cells, is a known causal gene in Charcot-Marie-Tooth disease type 1A and hereditary neuropathy with liability to pressure palsies (Sahenk et al., 1998; Watila & Balarabe, 2015). Chronic ethanol exposure suppressed *Pmp22* expression in female DRG and male spinal cord tissue, suggesting a convergent mechanism through which chronic ethanol disrupts myelin homeostasis in both sexes. Myelin- and axon-associated gene sets were enriched in all sex-tissue ethanol DEG sets, which validates these results given the well-characterized importance of these structures to peripheral neuropathies (Lehmann & Höke, 2010). Our laboratory has previously reported acute or chronic ethanol regulation of myelin-related genes in prefrontal cortex from mice, rhesus macaques and human autopsy material (Bogenpohl et al., 2019; Farris & Miles, 2013; Kerns et al., 2005; Lewohl et al., 2000; Wolstenholme et al., 2017). The effects of chronic ethanol on myelin regulation and myelin-producing oligodendrocytes have previously been reviewed by Rice and Gu (Rice & Gu, 2019) and Miguel-Hidalgo (Miguel-Hidalgo, 2018). The results here demonstrate that disruption of myelin gene expression by chronic ethanol may also be an important in the pathological mechanisms underlying AIPN.

Genes regulated by ethanol in these tissues were enriched for numerous neuroanatomical and ion channel GO terms. Myelin- and axon-associated gene sets were enriched in all sex-tissue ethanol DEG sets, which validates these results given the well-characterized importance of these structures to peripheral neuropathies (Lehmann & Höke, 2010). GO terms related to sodium ion transport were likewise enriched in male spinal cord, male DRG, and female DRG ethanol DEGs. Sodium channels regulate the firing of neurons, and Na+ channel-mediated neuron hyperexcitability is thought to be a basic mechanism of neuropathy (Devor, 2006). In the present study, two genes (*Scn1b* and *Scn4b*) were upregulated by ethanol in both male DRG and male spinal cord samples. Taken together, these GO enrichment results imply similar gene expression profiles between chronic ethanol and neuropathic pain.

To further contextualize our results, we compared our ethanol-regulated gene expression profiles to those identified in previous transcriptomic studies of peripheral neuropathy. Pokhilko et al. reported a set of genes differentially expressed across two rodent models and two surgical paradigms of neuropathic pain (Pokhilko et al., 2020). Despite differences in the underlying etiology, surgical injury versus ethanol-induced toxicity, we observed significant overlap between the Pokhilko DEG set and the ethanol-regulated genes in male DRG tissue. Among the overlapping genes were *Oprd1* (delta opioid receptor), *Sdc1* (syndecan-1), and *Cacna2d1* (a voltage-gated calcium channel auxiliary subunit). By their inclusion in the Pokhilko dataset, these genes have all been implicated in rodent neuropathic pain models. The convergence between our AIPN model and surgical models of neuropathy suggests that a core set of genes may underlie the molecular pathology of neuropathic pain, regardless of its origin. This overlap supports the existence of a shared “neuropathy signature” and highlights the potential translational relevance of our findings to broader pain research.

Sex differences in DRG-specific transcriptomic responses to surgically induced neuropathic pain have been documented previously (Liang et al., 2020; Lopes et al., 2017; Stephens et al., 2019). Mogil et al. meta-analyzed the results of these studies and found numerous GO categories enriched in sex-specific sets of neuropathic pain genes (Mogil et al., 2024). Many of these GO terms were similarly overrepresented in our ethanol DEG sets, including glial cell differentiation (GO:0010001; enriched in Male DRG, Male SC, Female DRG, Female SC EtOH DEGs), extracellular structure organization (GO:0043062, enriched in Female DRG, Female SC EtOH DEGs), sensory perception of pain (GO0019233, enriched in Male DRG, Female DRG EtOH DEGs), and potassium ion transmembrane transport (GO:0071805, enriched in Male DRG, Male SC EtOH DEGs). These convergent enrichment results again highlight cause-agnostic biological mechanisms that may be contributing to the development or persistence of peripheral neuropathy, and they underscore the importance of considering sex as a biological variable in preclinical pain and alcohol research.

To identify coordinated patterns of gene expression related to chronic ethanol exposure, we applied weighted gene co-expression network analysis (WGCNA) across each sex-tissue dataset. This method clusters genes into modules based on shared expression patterns, capturing functional relationships that may not be apparent through differential expression analysis alone. Of the resulting modules, 14 showed significant correlations between their eigengenes and ethanol treatment. GO enrichment analysis of these modules revealed consistent themes with our DEG results, including biological processes associated with axons, myelin structure, ion channels, and mitochondrial function. These findings further support the idea that ethanol disrupts cellular pathways critical to peripheral nerve integrity and neuronal excitability and structure.

WGCNA hub genes, highly interconnected module members, are strongly co-expressed with other genes in their corresponding modules. We identified hub genes for the 14 alcohol-associated WGCNA modules and conducted a search of the pain and neuropathy literature for these hubs. Strikingly, 10 of the 14 hubs (*Itga6*, *Map1lc3a*, *Cbln2*, *Pllp*, *Efhd1*, *Gas6*, *Srxn1*, *Jmjd1c*, *Klf9*, *Phf8*) were either the subject of pain-related studies or appeared in the results of pain-related experiments (Chang et al., 2018; Cui et al., 2022; Mamet et al., 2017; Ruan et al., 2024; Sandor et al., 2018; Sapio et al., 2021; Ulisse et al., 2020; Witucki & Jakubowski, 2023; Zhang et al., 2013; Zhang et al., 2025). For example, ozone therapy applied to a chronic constriction injury model of neuropathy in mice activated a *Gas6*-associated signaling pathway (Ruan et al., 2024). This pathway activated *Socs3*, yielding decreased neuroinflammation and reducing neuropathic pain. Co-inhibition of *Klf9*, a module hub gene, with *Klf6* and *Klf15* significantly attenuated mechanical hypersensitivity induced by spared nerve injury and chronic constriction injury in rats (Mamet et al., 2017). The hub gene *Itga6*, which codes for integrin alpha 6, was shown to partially underlie axonal regeneration following nerve injury through interactions with Schwann cells (Chang et al., 2018). *Phf8*, the hub for an alcohol-correlated module of male spinal cord genes, is a histone demethylase implicated in neuropathic pain via an autophagic pathway (Witucki & Jakubowski, 2023). Critically, hub *Jmjd1c* was recently identified as a member of a *Klf15/Jmjd1c*/*Socs3*/*JAK*/*STAT3* signaling pathway (Zhang et al., 2025). Downregulation of the *Jmjd1c* histone H3K9 demethylase attenuated *Socs3* expression, promoting the progression of neuropathic pain. *Jmjd1c* also regulates a pathway containing hub gene *Phf8* (GAS5-miR-495-3p-PHF8) in mouse cardiomyocytes, where silencing of *Jmjd1c* attenuated cardiac hypertrophy (Zhao et al., 2022). These hub genes offer potential mechanistic insight into ethanol-induced nerve damage and represent promising targets for therapeutic intervention in AIPN.

This study has several limitations that should be considered when interpreting the results. The female spinal cord dataset yielded relatively few differentially expressed genes, contrasting the robust DEG profiles observed in other sex-tissue groups. We hypothesize that this discrepancy may be due to inadvertent dissection variability, which could dilute transcriptional differences across experimental groups. Additionally, RNA samples from male and female mice were sequenced in separate cohorts, which may have introduced batch effects or confounded sex-specific transcriptional differences.

Future studies should incorporate mixed-sex cohorts to minimize potential biases. Finally, while the Lieber-DeCarli model simulates chronic high-dose alcohol exposure, it does not capture all aspects of alcohol use in humans (e.g., polysubstance use) which may also influence the development of peripheral neuropathy. These considerations underscore the need for replication in complementary models and careful tissue handling protocols in future transcriptomic investigations.

Alcohol-induced peripheral neuropathy affects millions of people around the world. With the sustained increase in alcohol consumption associated with the COVID-19 pandemic (Ayyala-Somayajula et al., 2025), AIPN is likely to produce an even greater burden on public health in the coming years. Our results recapitulate previous neuropathy transcriptomics work from surgical models of peripheral neuropathy, further validating the association of these genes with possible causal mechanisms of nerve degradation. Moreover, we identified key hub genes *Jmjd1c*, *Phf8*, *Gas6*, and *Itga6* as high-priority candidates for future investigation, given their regulatory roles in inflammation, axonal regeneration, and neuropathic pain signaling pathways. Future work should focus on validating the functional significance of these candidate genes in vivo and determining their relevance in human AIPN cohorts.

Figure S1. UpSet plot showing ethanol DEGs specific to, and shared across, sex-tissue groups. The number of genes in each set is indicated on top of the bars.

Figure S2. Heatmaps for each sex-tissue group showing correlations between WGCNA module eigengenes and ethanol consumption variables. We selected the binary EtOH variable to report due to the high concordance observed between this variable and the continuous measures. The number of genes per module is indicated in parentheses.

Table S1. A-D: Ethanol-regulated genes for each sex-tissue group as defined by an FDR < 0.1. E-F: Genes regulated in both tissues in male and female mice, respectively.

Table S2. Gene ontology enrichment results for all sex-tissue groups. Significant enrichment was defined as a false discovery rate (FDR) < 0.05, based on Benjamini-Hochberg-adjusted p-values.

Table S3. Overlapping genes between ethanol differentially expressed genes and the rodent neuropathy gene set reported by Table S2 in Pokhilko et al.

Table S4. Gene ontology enrichment results for WGCNA modules significantly correlated with ethanol exposure. Significant enrichment was defined as a false discovery rate (FDR) < 0.05, based on Benjamini-Hochberg-adjusted p-values.

Table S5. **All WGCNA modules with enrichment for corresponding sex-tissue group ethanol DEGs.**

Table S6. **Module membership for each gene detected in sex-tissue group-specific WGCNA.**

## References

Alford, D. P., German, J. S., Samet, J. H., Cheng, D. M., Lloyd-Travaglini, C. A., & Saitz, R. (2016). Primary Care Patients with Drug Use Report Chronic Pain and Self-Medicate with Alcohol and Other Drugs. J Gen Intern Med, 31(5), 486–491. 10.1007/s11606-016-3586-5

Ayyala-Somayajula, D., Dodge, J. L., Leventhal, A. M., Terrault, N. A., & Lee, B. P. (2025). Trends in Alcohol Use After the COVID-19 Pandemic: A National Cross-Sectional Study. Ann Intern Med, 178(1), 139–142. 10.7326/annals-24-02157

Barry, A. M., Zhao, N., Yang, X., Bennett, D. L., & Baskozos, G. (2023). Deep RNA-seq of male and female murine sensory neuron subtypes after nerve injury. Pain, 164(10), 2196–2215. 10.1097/j.pain.0000000000002934

Bogenpohl, J. W., Smith, M. L., Farris, S. P., Dumur, C. I., Lopez, M. F., Becker, H. C., Grant, K. A., & Miles, M. F. (2019). Cross-Species Co-analysis of Prefrontal Cortex Chronic Ethanol Transcriptome Responses in Mice and Monkeys. Front Mol Neurosci, 12, 197. 10.3389/fnmol.2019.00197

Bolívar, S., Sanz, E., Ovelleiro, D., Zochodne, D. W., & Udina, E. (2024). Neuron-specific RNA-sequencing reveals different responses in peripheral neurons after nerve injury. eLife, 12, RP91316. 10.7554/eLife.91316

Chang, I. A., Kim, K. J., & Namgung, U. (2018). α6 and β1 Integrin Heterodimer Mediates Schwann Cell Interactions with Axons and Facilitates Axonal Regeneration after Peripheral Nerve Injury. Neuroscience, 371, 49–59. 10.1016/j.neuroscience.2017.11.046

Chen, S., Zhou, Y., Chen, Y., & Gu, J. (2018). fastp: an ultra-fast all-in-one FASTQ preprocessor. Bioinformatics, 34(17), i884–i890. 10.1093/bioinformatics/bty560

Cui, C. Y., Liu, X., Peng, M. H., Liu, Q., & Zhang, Y. (2022). Identification of key candidate genes and biological pathways in neuropathic pain. Comput Biol Med, 150, 106135. 10.1016/j.compbiomed.2022.106135

Danecek, P., Bonfield, J. K., Liddle, J., Marshall, J., Ohan, V., Pollard, M. O., Whitwham, A., Keane, T., McCarthy, S. A., Davies, R. M., & Li, H. (2021). Twelve years of SAMtools and BCFtools. GigaScience, 10(2). 10.1093/gigascience/giab008

De Logu, F., Li Puma, S., Landini, L., Portelli, F., Innocenti, A., de Araujo, D. S. M., Janal, M. N., Patacchini, R., Bunnett, N. W., Geppetti, P., & Nassini, R. (2019). Schwann cells expressing nociceptive channel TRPA1 orchestrate ethanol-evoked neuropathic pain in mice. J Clin Invest, 129(12), 5424–5441. 10.1172/jci128022

Devor, M. (2006). Sodium Channels and Mechanisms of Neuropathic Pain. The Journal of Pain, 7(1), S3–S12. 10.1016/j.jpain.2005.09.006

Dobin, A., Davis, C. A., Schlesinger, F., Drenkow, J., Zaleski, C., Jha, S., Batut, P., Chaisson, M., & Gingeras, T. R. (2013). STAR: ultrafast universal RNA-seq aligner. Bioinformatics, 29(1), 15–21. 10.1093/bioinformatics/bts635

England, J. D., & Asbury, A. K. (2004). Peripheral neuropathy. The Lancet, 363(9427), 2151–2161. 10.1016/S0140-6736(04)16508-2

Farris, S. P., & Miles, M. F. (2013). Fyn-dependent gene networks in acute ethanol sensitivity. PLoS One, 8(11), e82435. 10.1371/journal.pone.0082435

Girach, A., Julian, T. H., Varrassi, G., Paladini, A., Vadalouka, A., & Zis, P. (2019). Quality of Life in Painful Peripheral Neuropathies: A Systematic Review. Pain Res Manag, 2019, 2091960. 10.1155/2019/2091960

Hinder, L. M., Murdock, B. J., Park, M., Bender, D. E., O’Brien, P. D., Rumora, A. E., Hur, J., & Feldman, E. L. (2018). Transcriptional networks of progressive diabetic peripheral neuropathy in the db/db mouse model of type 2 diabetes: An inflammatory story. Experimental Neurology, 305, 33–43. 10.1016/j.expneurol.2018.03.011

Julian, T., Glascow, N., Syeed, R., & Zis, P. (2019). Alcohol-related peripheral neuropathy: a systematic review and meta-analysis. J Neurol, 266(12), 2907–2919. 10.1007/s00415-018-9123-1

Kerns, R. T., Ravindranathan, A., Hassan, S., Cage, M. P., York, T., Sikela, J. M., Williams, R. W., & Miles, M. F. (2005). Ethanol-responsive brain region expression networks: implications for behavioral responses to acute ethanol in DBA/2J versus C57BL/6J mice. J Neurosci, 25(9), 2255–2266. 10.1523/jneurosci.4372-04.2005

Kokotis, P., Papantoniou, M., Schmelz, M., Buntziouka, C., Tzavellas, E., & Paparrigopoulos, T. (2023). Pure small fiber neuropathy in alcohol dependency detected by skin biopsy. Alcohol, 111, 67–73. 10.1016/j.alcohol.2023.05.006

Langfelder, P., & Horvath, S. (2008). WGCNA: an R package for weighted correlation network analysis. BMC Bioinformatics, 9(1), 559. 10.1186/1471-2105-9-559

Lehmann, H. C., & Höke, A. (2010). Schwann cells as a therapeutic target for peripheral neuropathies. CNS Neurol Disord Drug Targets, 9(6), 801–806. 10.2174/187152710793237412

Lewohl, J. M., Wang, L., Miles, M. F., Zhang, L., Dodd, P. R., & Harris, R. A. (2000). Gene expression in human alcoholism: microarray analysis of frontal cortex. Alcohol Clin Exp Res, 24(12), 1873–1882.

Li, Y., Yin, C., Liu, B., Nie, H., Wang, J., Zeng, D., Chen, R., He, X., Fang, J., Du, J., Liang, Y., Jiang, Y., Fang, J., & Liu, B. (2021). Transcriptome profiling of long noncoding RNAs and mRNAs in spinal cord of a rat model of paclitaxel-induced peripheral neuropathy identifies potential mechanisms mediating neuroinflammation and pain. Journal of Neuroinflammation, 18(1), 48. 10.1186/s12974-021-02098-y

Liang, Z., Hore, Z., Harley, P., Uchenna Stanley, F., Michrowska, A., Dahiya, M., La Russa, F., Jager, S. E., Villa-Hernandez, S., & Denk, F. (2020). A transcriptional toolbox for exploring peripheral neuroimmune interactions. Pain, 161(9), 2089–2106. 10.1097/j.pain.0000000000001914

Liao, Y., Smyth, G. K., & Shi, W. (2013). The Subread aligner: fast, accurate and scalable read mapping by seed-and-vote. Nucleic Acids Res, 41(10), e108. 10.1093/nar/gkt214

Lopes, D. M., Malek, N., Edye, M., Jager, S. B., McMurray, S., McMahon, S. B., & Denk, F. (2017). Sex differences in peripheral not central immune responses to pain-inducing injury. Scientific Reports, 7(1), 16460. 10.1038/s41598-017-16664-z

Love, M. I., Huber, W., & Anders, S. (2014). Moderated estimation of fold change and dispersion for RNA-seq data with DESeq2. Genome Biology, 15(12), 550. 10.1186/s13059-014-0550-8

Mamet, J., Klukinov, M., Harris, S., Manning, D. C., Xie, S., Pascual, C., Taylor, B. K., Donahue, R. R., & Yeomans, D. C. (2017). Intrathecal administration of AYX2 DNA-decoy produces a long-term pain treatment in rat models of chronic pain by inhibiting the KLF6, KLF9 and KLF15 transcription factors. Mol Pain, 13, 1744806917727917. 10.1177/1744806917727917

Miguel-Hidalgo, J. J. (2018). Molecular Neuropathology of Astrocytes and Oligodendrocytes in Alcohol Use Disorders. Front Mol Neurosci, 11, 78. 10.3389/fnmol.2018.00078

Mogil, J. S., Parisien, M., Esfahani, S. J., & Diatchenko, L. (2024). Sex differences in mechanisms of pain hypersensitivity. Neuroscience & Biobehavioral Reviews, 163, 105749. 10.1016/j.neubiorev.2024.105749

Moncayo, L. V., Adu-Gyamfi, F., Mohiuddin, A., Cruz, M., Dahman, A.-R., Akbar, Z., Kidd, S., Siddiqi, A., Khan, A., Chiang, K., Singh, T., Herz, S. M., Mawaldi, L., Valentia, G., Patel, P., Rauf, A., Miles, M. F., & Damaj, M. I. (2025). Peripheral Neuropathy After Chronic Alcohol Exposure in Mice: Impact of sex, total intake and duration and alcohol metabolism. bioRxiv, 2025.2009.2002.673579. 10.1101/2025.09.02.673579

Naratadam, G. T., Mecklenburg, J., Shein, S. A., Zou, Y., Lai, Z., Tumanov, A. V., Price, T. J., & Akopian, A. N. (2024). Degenerative and regenerative peripheral processes are associated with persistent painful chemotherapy-induced neuropathies in males and females. Scientific Reports, 14(1), 17543. 10.1038/s41598-024-68485-6

Pokhilko, A., Nash, A., & Cader, M. Z. (2020). Common transcriptional signatures of neuropathic pain. Pain, 161(7), 1542–1554. 10.1097/j.pain.0000000000001847

Rice, J., & Gu, C. (2019). Function and Mechanism of Myelin Regulation in Alcohol Abuse and Alcoholism. Bioessays, 41(7), e1800255. 10.1002/bies.201800255

Riley, J. L., 3rd, & King, C. (2009). Self-report of alcohol use for pain in a multi-ethnic community sample. J Pain, 10(9), 944–952. 10.1016/j.jpain.2009.03.005

Ruan, S., Jia, R., Hu, L., Liu, Y., Tian, Q., Jiang, K., Xia, X., Tao, X., Liu, W. T., Pan, Y., & Hu, F. (2024). Ozone promotes macrophage efferocytosis and alleviates neuropathic pain by activating the AMPK/Gas6-MerTK/SOCS3 signaling pathway. Front Immunol, 15, 1455771. 10.3389/fimmu.2024.1455771

Sahenk, Z., Chen, L., & Freimer, M. (1998). A novel *PMP22* point mutation causing HNPP phenotype. Neurology, 51(3), 702–707. doi:10.1212/WNL.51.3.702

Sandor, K., Krishnan, S., Agalave, N. M., Krock, E., Salcido, J. V., Fernandez-Zafra, T., Khoonsari, P. E., Svensson, C. I., & Kultima, K. (2018). Spinal injection of newly identified cerebellin-1 and cerebellin-2 peptides induce mechanical hypersensitivity in mice. Neuropeptides, 69, 53–59. 10.1016/j.npep.2018.04.004

Sanmaneechai, O., Feely, S., Scherer, S. S., Herrmann, D. N., Burns, J., Muntoni, F., Li, J., Siskind, C. E., Day, J. W., Laura, M., Sumner, C. J., Lloyd, T. E., Ramchandren, S., Shy, R. R., Grider, T., Bacon, C., Finkel, R. S., Yum, S. W., Moroni, I., . . . Consortium, f. t. I. N. C.-R. D. C. R. (2015). Genotype–phenotype characteristics and baseline natural history of heritable neuropathies caused by mutations in the MPZ gene. Brain, 138(11), 3180–3192. 10.1093/brain/awv241

Sapio, M. R., Kim, J. J., Loydpierson, A. J., Maric, D., Goto, T., Vazquez, F. A., Dougherty, M. K., Narasimhan, R., Muhly, W. T., Iadarola, M. J., & Mannes, A. J. (2021). The Persistent Pain Transcriptome: Identification of Cells and Molecules Activated by Hyperalgesia. J Pain, 22(10), 1146–1179. 10.1016/j.jpain.2021.03.155

Shen, L. (2025). GeneOverlap: Test and visualize gene overlaps. In (R package version 1.44.0 ed.). Icahn School of Medicine at Mount Sinai.

Signal, B., & Kahlke, T. (2022). how_are_we_stranded_here: quick determination of RNA-Seq strandedness. BMC Bioinformatics, 23(1), 49. 10.1186/s12859-022-04572-7

Stephens, K. E., Zhou, W., Ji, Z., Chen, Z., He, S., Ji, H., Guan, Y., & Taverna, S. D. (2019). Sex differences in gene regulation in the dorsal root ganglion after nerve injury. BMC Genomics, 20(1), 147. 10.1186/s12864-019-5512-9

Tessitore, M. E., Pereira-Rufino, L. d. S., Panfilio, C. E., de Cassia Sinigaglia, R., Júnior, O. A., Maluf, L. L.-S., Conte, R., Ladd, F. V. L., & Céspedes, I. C. (2022). Alcoholic neuropathy associated with chronic alcohol intake. IBRO Neuroscience Reports, 13, 177–186. 10.1016/j.ibneur.2022.08.004

Thompson, T., Oram, C., Correll, C. U., Tsermentseli, S., & Stubbs, B. (2017). Analgesic Effects of Alcohol: A Systematic Review and Meta-Analysis of Controlled Experimental Studies in Healthy Participants. J Pain, 18(5), 499–510. 10.1016/j.jpain.2016.11.009

Ulisse, V., Dey, S., Rothbard, D. E., Zeevi, E., Gokhman, I., Dadosh, T., Minis, A., & Yaron, A. (2020). Regulation of axonal morphogenesis by the mitochondrial protein Efhd1. Life Sci Alliance, 3(7). 10.26508/lsa.202000753

Visser, N. A., Notermans, N. C., Linssen, R. S., van den Berg, L. H., & Vrancken, A. F. (2015). Incidence of polyneuropathy in Utrecht, the Netherlands. Neurology, 84(3), 259–264. 10.1212/wnl.0000000000001160

Watila, M. M., & Balarabe, S. A. (2015). Molecular and clinical features of inherited neuropathies due to PMP22 duplication. J Neurol Sci, 355(1-2), 18–24. 10.1016/j.jns.2015.05.037

Welleford, A. S., Quintero, J. E., Seblani, N. E., Blalock, E., Gunewardena, S., Shapiro, S. M., Riordan, S. M., Huettl, P., Guduru, Z., Stanford, J. A., van Horne, C. G., & Gerhardt, G. A. (2020). RNA Sequencing of Human Peripheral Nerve in Response to Injury: Distinctive Analysis of the Nerve Repair Pathways. Cell Transplant, 29, 963689720926157. 10.1177/0963689720926157

Wickham, H. (2016). ggplot2 : elegant graphics for data analysis (2nd edition ed.). Springer International Publishingr.

Witucki, Ł., & Jakubowski, H. (2023). Homocysteine metabolites inhibit autophagy, elevate amyloid beta, and induce neuropathy by impairing Phf8/H4K20me1-dependent epigenetic regulation of mTOR in cystathionine β-synthase-deficient mice. J Inherit Metab Dis, 46(6), 1114–1130. 10.1002/jimd.12661

Wöhrle, J. C., Spengos, K., Steinke, W., Goebel, H. H., & Hennerici, M. (1998). Alcohol-Related Acute Axonal Polyneuropathy: A Differential Diagnosis of Guillain-Barré Syndrome. Archives of Neurology, 55(10), 1329–1334. 10.1001/archneur.55.10.1329

Wolstenholme, J. T., Mahmood, T., Harris, G. M., Abbas, S., & Miles, M. F. (2017). Intermittent Ethanol during Adolescence Leads to Lasting Behavioral Changes in Adulthood and Alters Gene Expression and Histone Methylation in the PFC. Front Mol Neurosci, 10, 307. 10.3389/fnmol.2017.00307

Yu, G., Wang, L. G., Han, Y., & He, Q. Y. (2012). clusterProfiler: an R package for comparing biological themes among gene clusters. Omics, 16(5), 284–287. 10.1089/omi.2011.0118

Zhang, E., Yi, M. H., Ko, Y., Kim, H. W., Seo, J. H., Lee, Y. H., Lee, W., & Kim, D. W. (2013). Expression of LC3 and Beclin 1 in the spinal dorsal horn following spinal nerve ligation-induced neuropathic pain. Brain Res, 1519, 31–39. 10.1016/j.brainres.2013.04.055

Zhang, L., Xie, Y., Wang, S., Gong, M., Chen, Z., Wang, C., & Li, P. (2025). Enhancer profiling uncovers Jmjd1c as an essential suppressor in neuropathic pain by targeting Socs3. Genes & Diseases, 101545. 10.1016/j.gendis.2025.101545

Zhao, L., Qi, F., Du, D., & Wu, N. (2022). Histone demethylase KDM3C regulates the lncRNA GAS5–miR-495-3p–PHF8 axis in cardiac hypertrophy. Annals of the New York Academy of Sciences, 1516(1), 286–299. 10.1111/nyas.14813

