## Supplementary figures and images for "Transcriptomic and network analyses of an alcohol-induced peripheral neuropathy model identify putative role for histone demethylase *Jmjd1c*"

### Figure S1

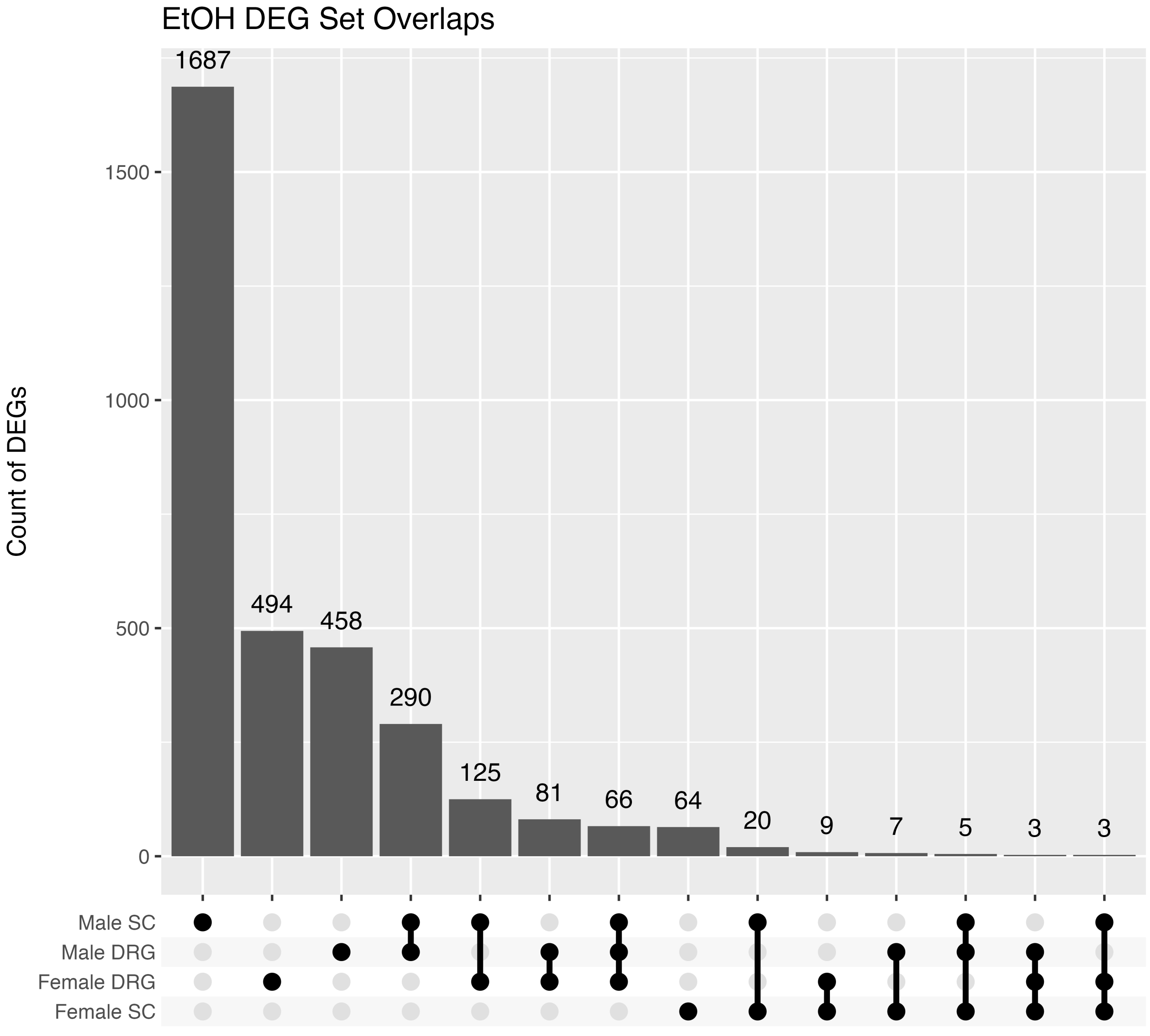

### Figure S2

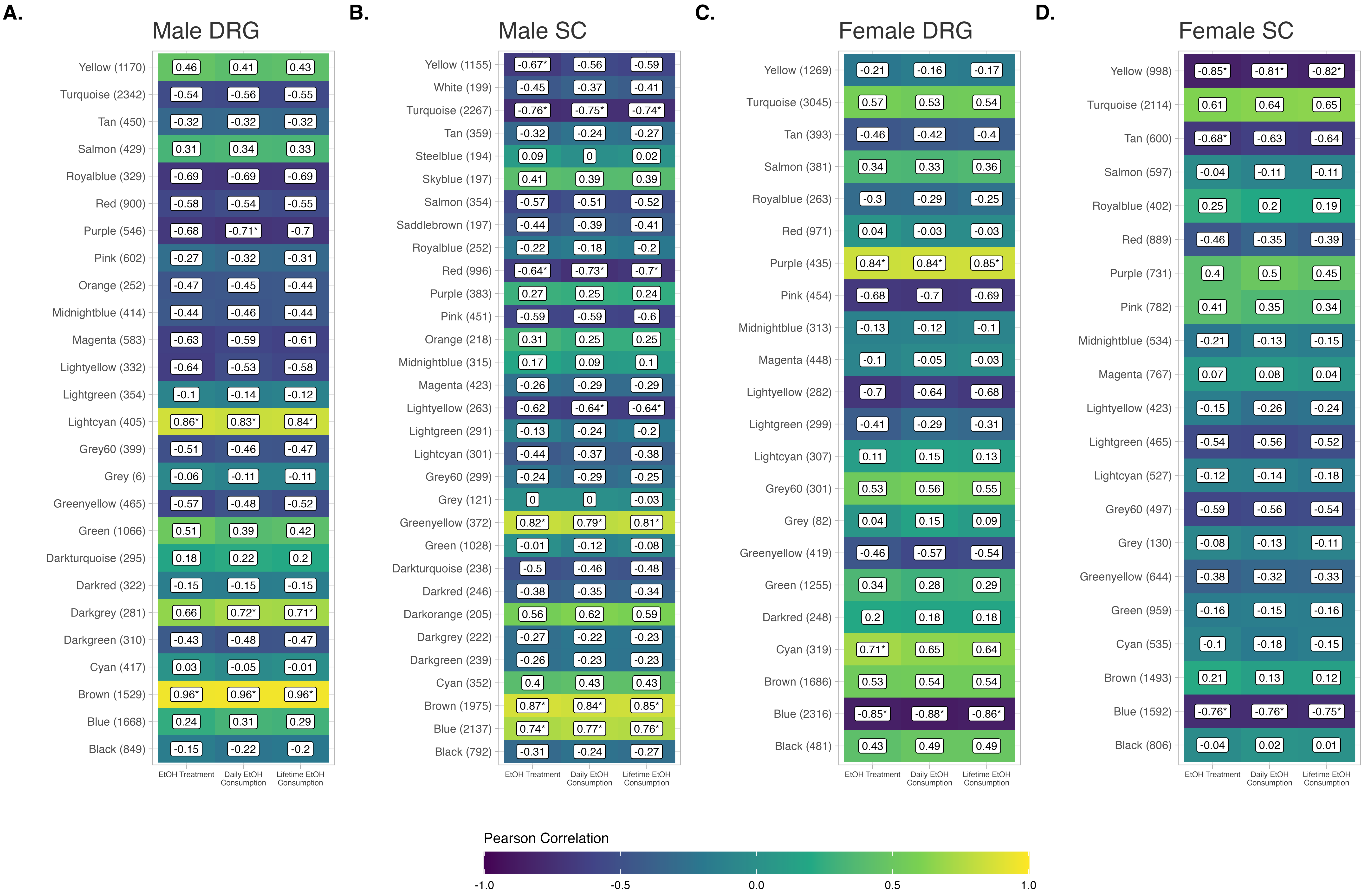
